# Single-cell landscape of bone marrow metastases in human neuroblastoma unraveled by deep multiplex imaging

**DOI:** 10.1101/2020.09.30.321539

**Authors:** Daria Lazic, Florian Kromp, Michael Kirr, Filip Mivalt, Fikret Rifatbegovic, Florian Halbritter, Marie Bernkopf, Andrea Bileck, Marek Ussowicz, Inge M Ambros, Peter F Ambros, Christopher Gerner, Ruth Ladenstein, Christian Ostalecki, Sabine Taschner-Mandl

**Affiliations:** St. Anna Children’s Cancer Research Institute (CCRI), Vienna, Austria; Department of Dermatology, University Hospital Erlangen, Erlangen, Germany; Department of Analytical Chemistry, Faculty of Chemistry, University of Vienna, Vienna, Austria; Department and Clinic of Pediatric Oncology, Hematology and Bone Marrow Transplantation, Wroclaw Medical University

## Abstract

Bone marrow commonly serves as a metastatic niche for disseminated tumor cells (DTCs) of solid cancers in patients with unfavorable clinical outcome. Single-cell assessment of bone marrow metastases is essential to decipher the entire spectrum of tumor heterogeneity in these cancers, however, has previously not been performed.

Here we used multi-epitope-ligand cartography (MELC) to spatially profile 20 biomarkers and assess morphology in DTCs as well as hematopoietic and mesenchymal cells of eight bone marrow metastases from neuroblastoma patients. We developed DeepFLEX, a single-cell image analysis pipeline for MELC data that combines deep learning-based cell and nucleus segmentation and overcomes frequent challenges of multiplex imaging methods including autofluorescence and unspecific antibody binding.

Using DeepFLEX, we built a single-cell atlas of bone marrow metastases comprising more than 35,000 single cells. Comparisons of cell type proportions between samples indicated that microenvironmental changes in the metastatic bone marrow are associated with tumor cell infiltration and therapy response. Hierarchical clustering of DTCs revealed multiple phenotypes with highly diverse expression of markers such as FAIM2, an inhibitory protein in the Fas apoptotic pathway, which we propose as a complementary marker to capture DTC heterogeneity in neuroblastoma.

The presented single-cell atlas provides first insights into the heterogeneity of human bone marrow metastases and is an important step towards a deeper understanding of DTCs and their interactions with the bone marrow niche.

## INTRODUCTION

Metastasis is the major cause of cancer-related deaths^1^ and relies on the ability of tumor cells to disseminate from the primary site and adapt to distant tissue environments.^2^ This is an arduous process, which can fuel heterogeneity among metastasizing and disseminated tumor cells (DTCs).^3^ Tumor heterogeneity manifests in variations of clinically important features such as the abundance of prognostic markers as well as therapeutic targets, which complicates patient stratification and explains failure of therapeutic approaches.^4,5,6,7^

Cancer cells are attracted by distant microenvironments that promote their growth and survival.^8^ One such hospitable microenvironment is the bone marrow, which has a major role in dormancy and relapse^9^ and is a frequent site of dissemination in numerous solid cancers^10,11^, such as breast cancer, colorectal cancer and neuroblastoma.^12^

Neuroblastoma, an extracranial neoplasm of the sympathetic nervous system, is the most common solid tumor in children in their first year of life and accounts for roughly 15% of childhood cancer related deaths.^13,14,15^ In more than 90% of metastatic stage (stage M) neuroblastoma patients, tumor cells disseminate to the bone marrow^16,17^, where some tumor cells may resist initial chemotherapy and give rise to relapse. These relapse seeding clones are frequently, already at the time-point of diagnosis, detected in the bone marrow, but not in the primary tumor.^6^ Based on bulk RNA-sequencing (RNA-seq), we have previously shown differences between the transcriptome of DTCs with predominantly hypoxia-associated genes enriched, and primary tumor cells with an increased expression of mesenchymal genes.^18^ Subsequently, two studies of neuroblastoma cell lines and primary tumors unraveled the gene regulatory networks driving two plastic phenotypes, adrenergic and mesenchymal type neuroblastoma cells and highlighted their importance, as the latter were more frequently found in post-therapy and relapse samples and were more resistant to chemotherapy.^19,20^ Thus, genetic and phenotypic tumor heterogeneity can be considered key to why treatment of metastatic disease remains poor.

Although phenotypic tumor heterogeneity of solid cancers has been investigated at the primary site at single-cell resolution,^21,22,23,24^ to date no analyses of human bone marrow metastases have been undertaken.^25^

In recent years, numerous technologies for the analysis of single cells have emerged and advanced rapidly. While single-cell RNA-seq (scRNA-seq) methods^26^ enable high-dimensional analyses of cells at the transcriptomic level, highly multiplexed imaging methods^27^ provide an image of every cell and thereby allow subcellular localization of proteins as well as morphological assessment. Despite the volume of developing multiplex imaging methods, the standard method to detect DTCs in bone marrow aspirates in neuroblastoma routine diagnostic procedures, is still automated immunofluorescence plus fluorescence in situ hybridization (AIPF), which is limited to only three biomarkers (GD2, CD56 and one genetic marker).^28,29^ In order to unravel the complete scope of intra-tumor heterogeneity and capture therapy-related changes and resistant cells in solid cancers with bone marrow involvement, a comprehensive single-cell map of bone marrow metastases is indispensable. Thus, we sought to provide the first single-cell atlas of bone marrow metastases including DTCs and cells of the microenvironment, by employing neuroblastoma as a model.

We applied Multi-Epitope-Ligand Cartography (MELC), a multiplex imaging method with a resolution of 450nm that employs automated sequential cycles of staining with fluorophore-coupled antibodies followed by immunofluorescence (IF) microscopy and photobleaching.^30,31^ A 20-plex antibody panel was established, and we developed an image analysis pipeline, called DeepFLEX, which tackles frequent obstacles of IF-based imaging and in addition involves accurate, deep learning-based cell and nucleus segmentation. Our study revealed novel markers, including FAIM2 (Fas Apoptotic Inhibitory Molecule 2), to capture heterogeneity of DTCs in bone marrow metastases of neuroblastoma patients. Moreover, our analyses delivered the first indication that the presence of DTCs as well as treatment are associated with dynamic changes in the bone marrow microenvironment.

## RESULTS

### Comprehensive single-cell multiplex immunofluorescence imaging panel

To analyze bone marrow metastases on a single-cell level, we sought to establish a MELC panel specific to neuroblastoma DTCs, hematopoietic, and mesenchymal cells in the bone marrow. Therefore, in our workflow (Fig. 1a), we first selected DTC-associated biomarkers, which we then validated separately by conventional IF staining.

**Figure 1:**
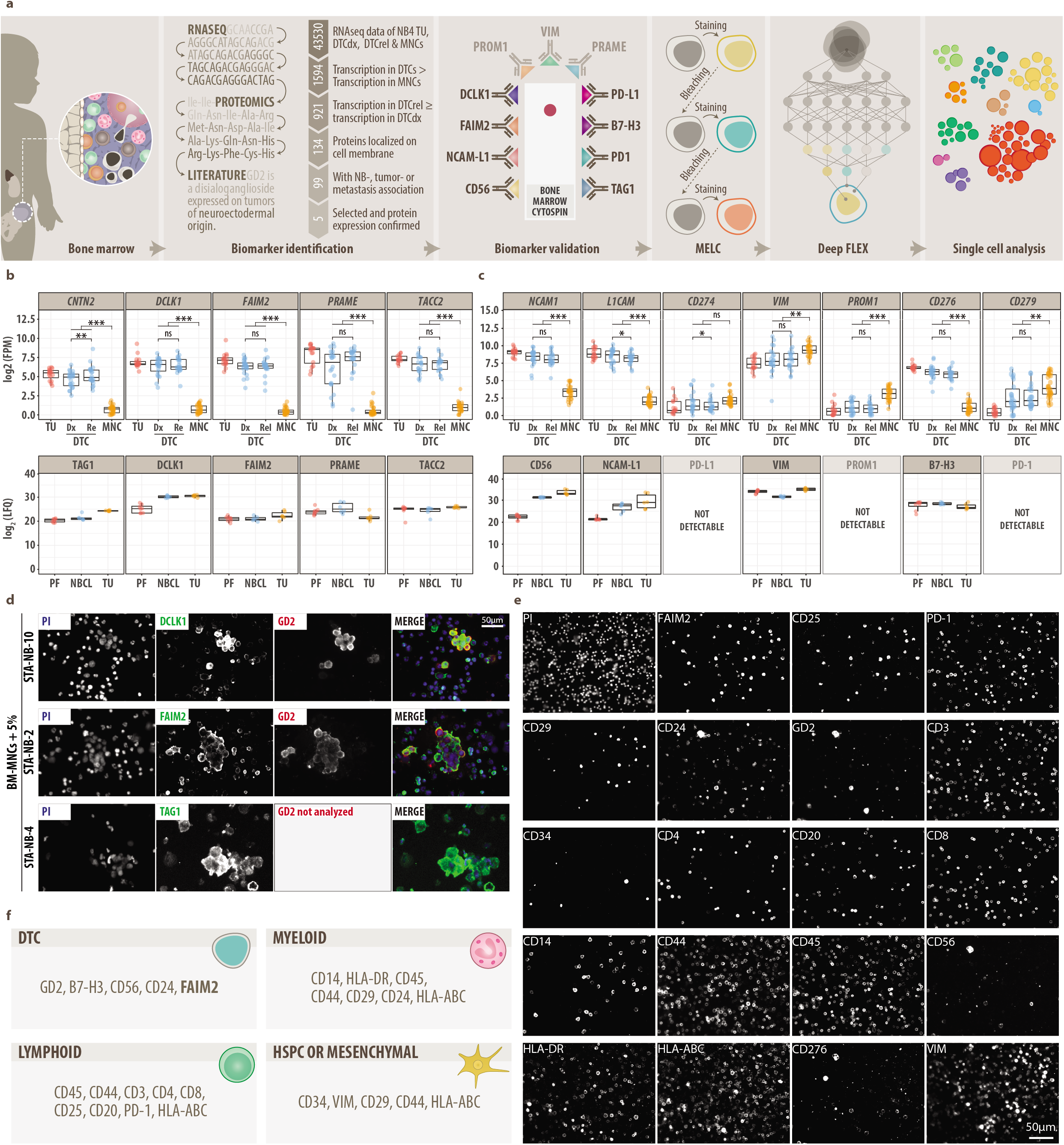
Data mining and establishment of multiplex imaging panel (MELC) **a,** Flow chart of experimental approach. **b,** 5 potential disseminated tumor cell (DTC) biomarkers identified by data mining of RNA-seq data, proteomics data (LC-MS/MS) and literature. Top: mRNA transcription (RNA-seq) in neuroblastoma primary tumors (TU), diagnostic (dX) and relapse (rel) DTCs and bone marrow-derived mononuclear cells (BM-MNCs). DESeq2, FDR-adjusted p value: ns, p> 0.05, *, p ≤ 0.05; **, p ≤ 0.01; ***, p ≤ 0.001. Bottom: Protein expression in NB peripheral-nerve-associated fibroblasts (PF), neuroblastoma cells lines and TU samples. LFQ, Label Free Quantification; FPM, fragments per million. **c,** Extension of potential DTC biomarkers by immune checkpoint molecules (PD-L1, PD-1, B7-H3), mesenchymal-type neuroblastoma cell markers (VIM, PROM1), therapeutic target NCAM-L1 and diagnostic neuroblastoma marker CD56. DESeq2, FDR-adjusted p value: ns, p> 0.05, *, p ≤ 0.05; **, p ≤ 0.01; ***, p ≤ 0.001. **d,** Representative MELC images of newly identified DTC biomarkers DCLK1, FAIM2 and TAG1 on separate samples stained by MELC. Top: DCLK1 (green) and GD2 (red) on BM-MNCs and neuroblastoma cells line STA-NB-10 (mixed 20:1); center: FAIM2 (green) and GD2 (red) on BM-MNCs and neuroblastoma cell line STA-NB-2 (20:1), bottom: TAG1 (green) on peripheral blood-derived MNCs and neuroblastoma cell line STA-NB-4 (20:1) stimulated with IFNγ and anti-CD3/28 beads. Nuclei were counterstained with DAPI (blue). **e,** Representative MELC images of our single-cell 20-plex panel on one patient bone marrow sample. **f,** Single-cell 20-plex panel composed of DTC, myeloid, lymphoid, mesenchymal and HSPC (hematopoietic stem and progenitor cell) markers.

Data mining (see Methods) based on RNA-seq data of stage M neuroblastoma primary tumors, DTCs, and bone marrow-derived mononuclear cells (MNCs); proteomics data of neuroblastoma tumors, neuroblastoma cell lines, and peripheral-nerve-associated fibroblasts; and public databases (Uniprot, Protein Atlas, PubMed) revealed five potential DTC biomarkers: TAG1 (*CNTN2*), DCLK1, FAIM2, PRAME and TACC2 (Fig. 1b). All five candidates had (I) significantly higher transcript levels in DTCs as compared to bone-marrow-derived MNCs, (II) an equal or higher transcription at the time point of relapse as compared to diagnosis, (III) literature available on neuroblastoma-, tumor- or metastasis association and (IV) protein expression described to be localized on the cell membrane and detected by proteomic analysis. To further characterize DTC heterogeneity, we complemented these five markers by mesenchymal-type neuroblastoma cell markers^20^ VIM and PROM1, immune checkpoint molecules B7-H3^32^ and PD-L1 with its binding partner PD-1^33^, the currently investigated therapeutic target NCAM-L1^34^, as well as two gold-standard neuroblastoma markers CD56 and the ganglioside GD2.^35^ VIM was highly, but not exclusively expressed by DTCs (Fig. 1c, top). PD-1, PD-L1 and PROM1 showed low expression on the mRNA level and were not detected in the mass spectrometry data (Fig. 1c, bottom), which can be explained by their expression on rare cell types.

To assess their protein expression on a single-cell level, we validated these 13 DTC-related biomarkers (Fig. S1a) using optimized IF-sample preparation protocols (Fig. S1b and S2a, Table S1) on neuroblastoma cell lines in isolation (Fig. S1c, Table S2 and S3) and spiked into bone marrow-derived MNCs or peripheral blood-derived MNCs (Fig. S2b). Three out of the 5 initial DTC biomarker candidates (Fig. 1b) yielded a tumor-specific IF-staining on neuroblastoma cell lines spiked into bone marrow-derived MNCs (Fig. 1d). These were DCLK1, which is crucial for neuroblast proliferation^36,37^, TAG1, a promoter of glioma proliferation^38^ and the inhibitor of Fas induced apoptosis FAIM2 (Fig. S3). We selected FAIM2 together with GD2, CD56, VIM, B7-H3 and PD-1 for multiplex imaging and tested them simultaneously with 14 other bone marrow hematopoietic and mesenchymal cell markers (Table S4) in MELC assays, finally resulting in a specific and robust 20-plex panel (Fig. 1e, Table 1).

**Table 1.**
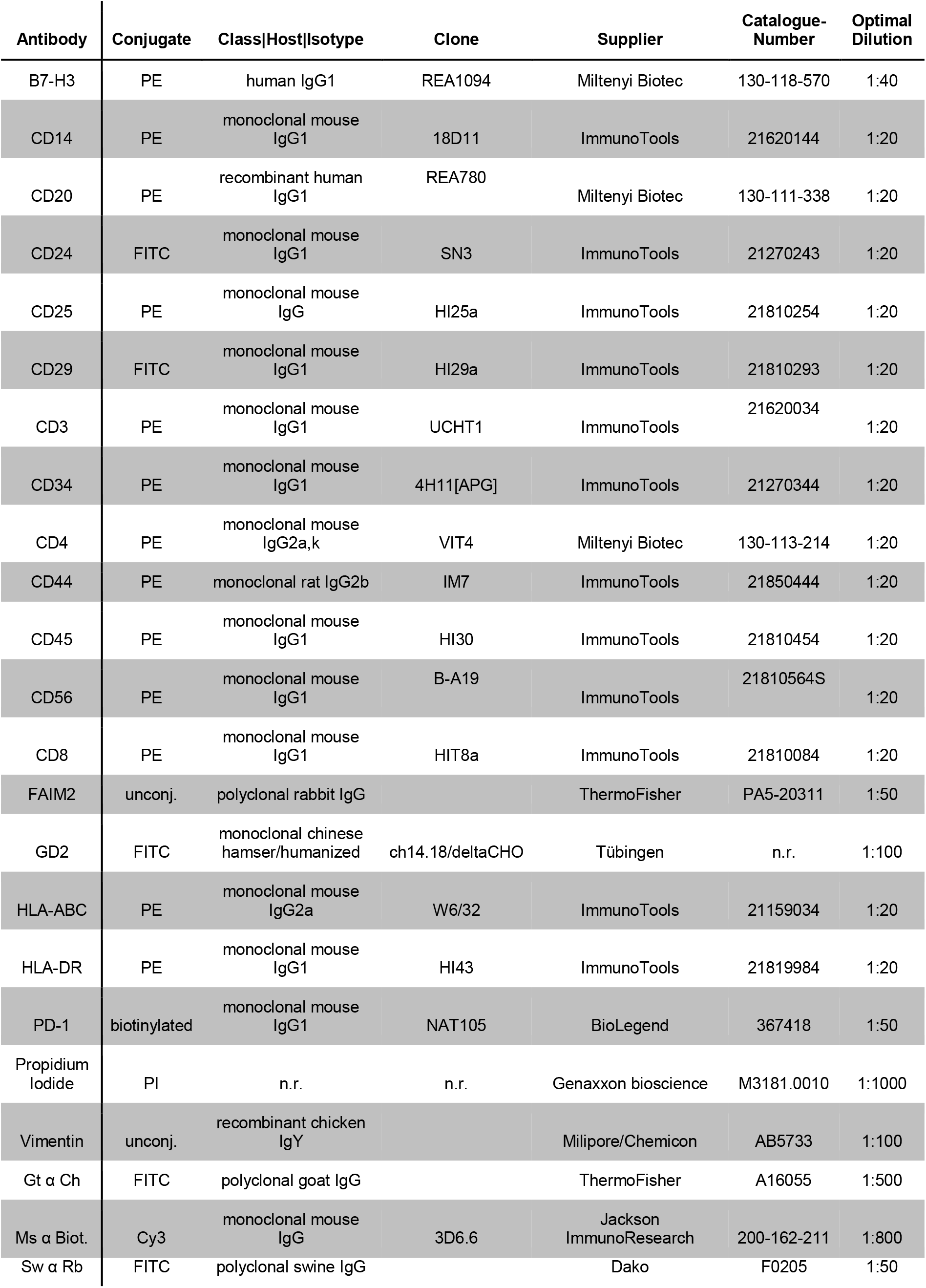
All primary and secondary antibodies, which passed the validation procedure and were included in the final 20-plex

In conclusion, we here provide a validated 20-plex MELC panel for neuroblastoma composed of DTC markers, including a novel candidate marker called FAIM2, as well as myeloid, lymphoid, mesenchymal, and hematopoietic stem and progenitor cell markers (Fig. 1f).

### Deep learning-based single-cell analysis pipeline for FLuorescence multiplEX imaging – DeepFLEX

MELC and other IF-based multiplex imaging methods suffer from inhomogeneous illumination, background noise due to incomplete signal removal by photobleaching or heat denaturation, autofluorescence and unspecific binding, which are either not addressed or not effectively solved in published single-cell image analysis pipelines.^39,40,41,42,43,44^

To address these challenges and allow unsupervised single-cell analysis of MELC imaging data, we developed DeepFLEX (Fig. 2a), a semi-automated, deep learning-based pipeline. DeepFLEX integrates methods for image processing, segmentation, feature extraction, normalization, and single-cell analysis that were recently published by our group and experts in the field (see Methods).

**Figure 2:**
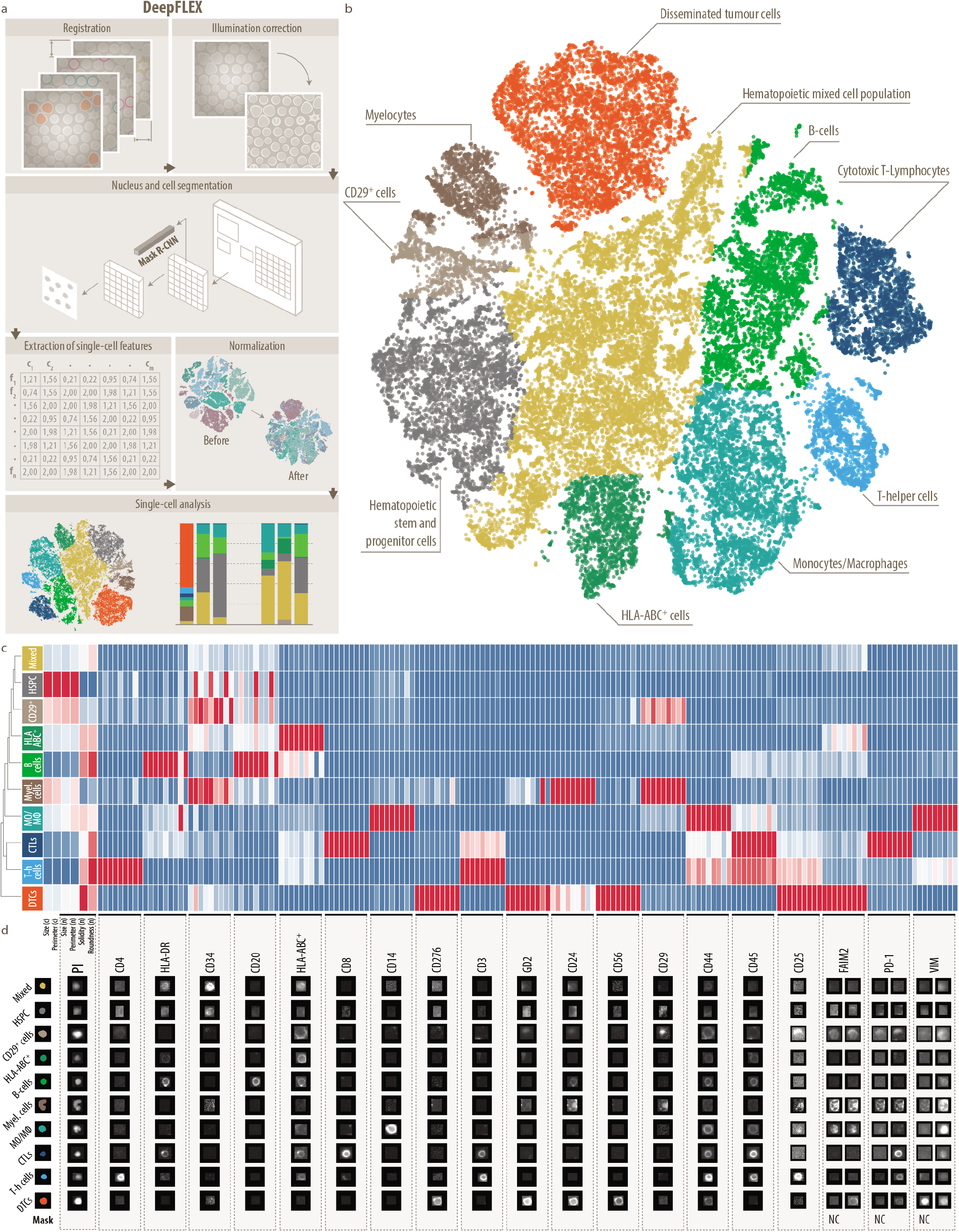
DeepFLEX and single-cell map of 8 bone marrow metastases from four patients with stage M and Ms neuroblastoma. **a,** Schematic overview of the deep learning-based image processing, segmentation, normalization and single-cell analysis pipeline DeepFLEX. **b,** Single-cell atlas of 35,700 single cells clustered colored by cell type. Dimensionality reduction was performed by A-tSNE (approximated and user steerable t-distributed Stochastic Neighbor Embedding) and subsequent clustering by GMS (Gaussian Mean Shift) in Cytosplore.^50^ **c,** Heatmap showing the median feature expression of all created clusters with feature-wise scaling. n, nucleus; c, cell. 9 columns per marker represent, from left to right, mean intensity, total intensity and mean of the highest 20% of pixel values in the (I) nucleus, (II) cell and (III) cytoplasm/membrane. DTCs, disseminated tumor cells; Myel., myelocytes; MO/MΦ, monocytes/macrophages; HSPC, hematopoietic stem and progenitor cells; T-h cells, T-helper cells; CTLs; cytotoxic T-lymphocytes; Mixed, hematopoietic mixed cell population. **d,** Representative gallery images of all cell types. For FAIM2, PD-1 and VIM we introduced negative controls (NC) to be used for normalization during data processing. Hence, for these four biomarkers, the ratio between right column and left column (NC) represents the true signal.

DeepFLEX corrects for common problems of IF staining and microscopy. Cross-correlation-based registration^45^ aligns images, which are shifted due to the microscope stage movement in between staining and bleaching cycles. Flat-field correction eliminates gross variations in illumination using calibration images. Image subtraction of the bleached from the subsequently stained sample removes bleaching remnants. CIDRE^46^, a retrospective multi-image illumination correction method based on energy minimization further homogenizes illumination towards image borders (Fig. S4a-d).

The deep neuronal network Mask R-CNN^47^, trained on an annotated fluorescence image dataset^48^, allows accurate cell and nucleus segmentation in the processed images. Simultaneous segmentation of the nucleus (based on the nuclear stain propidium iodide) and the cell (based on phase contrast images acquired prior to each staining cycle) allows the elimination of displaced and inaccurately segmented cells by considering only those cells for the analysis, which are present in every cell segmentation mask, but also in the nucleus segmentation mask. Cells affected by artifacts are excluded by user-guided region selection (Fig. S4e-h).

Based on the cell and the nucleus segmentation mask, DeepFLEX facilitates the extraction of single-cell features (Table S5) describing the intensity (mean intensity, total intensity, and mean of the highest 20% of pixel values) and morphology (roundness, solidity, perimeter and size) of three cell compartments, namely the cell itself, the nucleus, and their difference, which represents the cell cytoplasm with the membrane (Fig. S5a-b).

Our pipeline diminishes non-specific staining, autofluorescence, and experimental batch effects by normalizing single-cell features based on images of negative control secondary antibodies (see Methods), and mutually exclusive marker pairs for the prediction of background levels via RESTORE^49^ (Table S6, Fig. S5c-d).

DeepFLEX analyzes the normalized single-cell data based on the integration of the visual analysis framework Cytosplore^50^ and the python data visualization library *seaborn* (seaborn.pydata.org), which provide methods for interactive and quantitative analysis of individual cell classes, respectively (Fig. S5e).

We proved that features extracted by DeepFLEX are comparable to their respective images in representing cells, by using them as inputs to shallow and deep neural networks trained on an annotated cell dataset and inferring cell classes (see Suppl. Methods, Fig. S6a-d, Table S7). In summary, DeepFLEX represents a comprehensive single-cell image analysis pipeline for MELC multiplex imaging, which includes accurate deep learning-based cell and nucleus segmentation, and demonstrates computational solutions for common obstacles of targeted multiplex imaging technologies such as unspecific binding and autofluorescence

### Single-cell map of tumor cells and the microenvironment in neuroblastoma bone marrow metastases

To obtain a single-cell map of DTCs and the bone marrow microenvironment in children with neuroblastoma, we used the designed 20-plex panel for MELC imaging of eight bone marrow samples (Table 2) collected from three stage M and one stage Ms neuroblastoma patient (Table 3) at different time-points during therapy. We then fed the generated multiplex images into DeepFLEX, which resulted in an atlas of 35,700 single cells distributed between ten clusters (Fig. 2b).

**Table 2.**
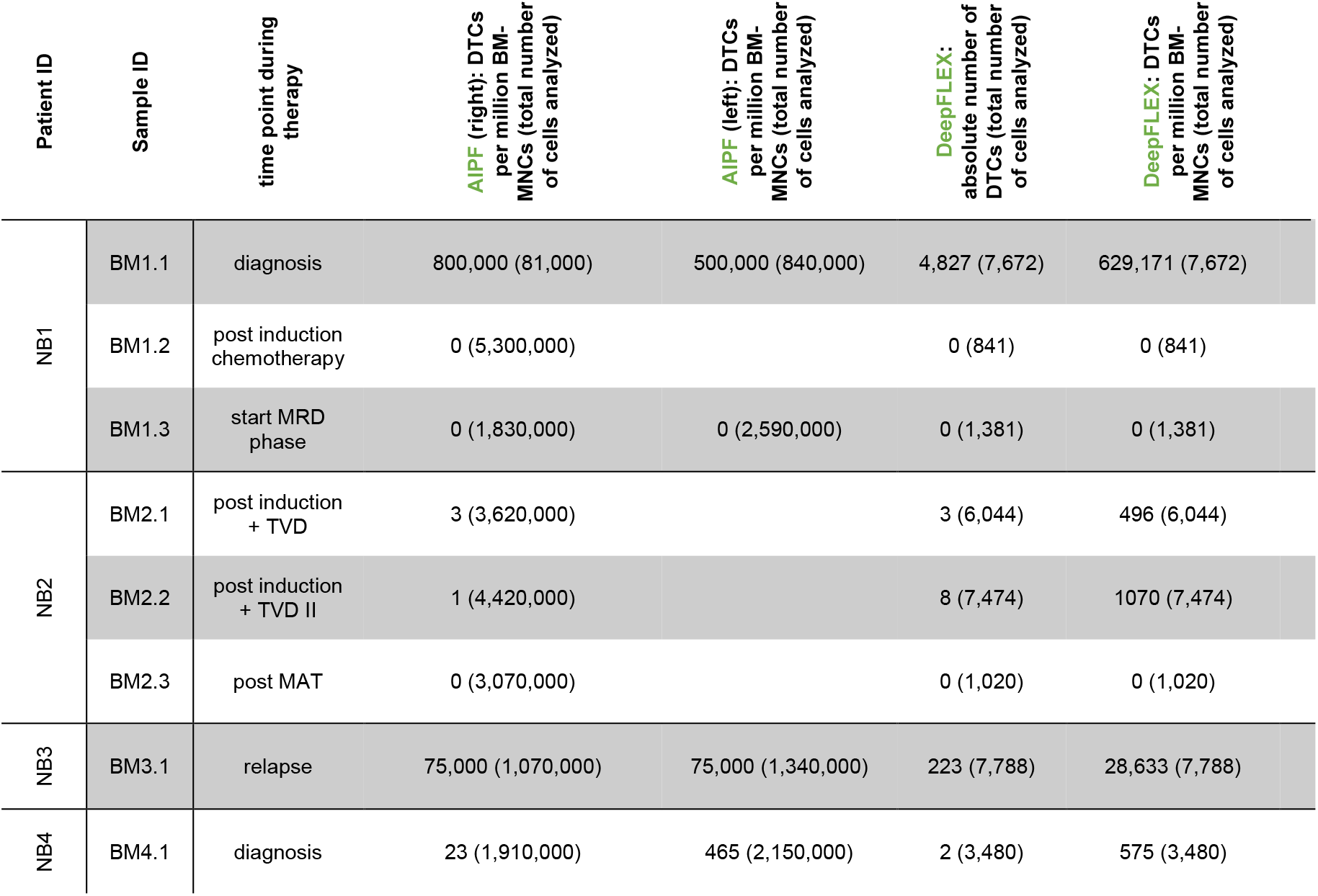
Sample set comprised of eight bone marrow samples. Tumor cell content detected by DeepFLEX was compared to AIPF, the standard method used in the diagnostic routine. AIPF, Automatic Immunofluorescence Plus FISH; BM-MNCs, bone marrow-derived mononuclear cells; left/right, bone marrow aspirate from left/right puncture side (pooled for analysis by DeepFLEX); TVD, topotecan-vincristine-doxorubicin; MAT, myeloablative therapy with autologous stem cell transplantation.

**Table 3.** Patient set. INRG, International Neuroblastoma Risk Group Staging System, HRNBL1, High Risk Neuroblastoma study 1; LINES, Low and Intermediate Risk Neuroblastoma European Study; MNA, *MYCN* amplification; neg, negative; pos, positive;

| Patient_ID | sex | age at diagnosis (months) | INRG | clinical study | event | dead/alive | MNA | 17q gain |
| --- | --- | --- | --- | --- | --- | --- | --- | --- |
| NB1 | F | 54 | M | HRNBL1 | no | alive | neg | pos |
| NB2 | M | 8.5 | M | no | no | alive | neg | pos |
| NB3 | F | 86 | M | HRNBL1 | yes | dead | pos | neg |
| NB4 | F | 5 | Ms | LINES | no | alive | neg | neg |

After confirming that potential batch effects have been eliminated (Fig. S5d, bottom), we annotated clusters based on median feature expression of cell-type-specific marker proteins (Fig. 2c, Fig. S7 and S8) and a recently published single-cell atlas^51^ of healthy adult human bone marrow. In addition, we verified our annotation using representative gallery images of each cell type (Fig. 2d).

We found most of the expected immune cell types including T-helper cells (T-h cells), cytotoxic T-lymphocytes (CTLs), monocytes and macrophages (MO/MΦ) as well as B-cells. A dominant proportion of cells in the bone marrow microenvironment of children represented a hematopoietic mixed (Fig. 2b, yellow cluster) and a stem and progenitor (Fig. 2b, grey cluster) cell phenotype. Moreover, we were able to identify a segregated tumor cell cluster with co-expression of all markers expressed by DTCs from our panel (GD2, CD56, B7-H3, CD24 and FAIM2). The mesenchymal marker VIM showed the highest expression on monocytes and macrophages (Fig. 2c, Fig. S8). In bulk transcriptomic and proteomic data (Fig 1c), VIM was also highly expressed in neuroblastoma cells, which was in accordance with the IF staining results on neuroblastoma cell lines (Fig. S1c, Fig. S2b). However, in the eight analyzed bone marrow samples, which were prepared with the same protocol (Fig. S2a) and stained with the identical antibody (Table 1), DTCs were negative for VIM (Fig. 2c, Fig. S8), and also the mesenchymal marker CD29, indicating that DTCs in our sample set are predominantly of an adrenergic type, but clarification will require further robust mesenchymal markers. CD29 was enriched in two clusters (Fig. 2c): (I) in myelocytes, which, in accordance with a previous study^52^, also showed a strong expression of CD24 as well as a c-shaped morphology (Fig. 2d), and hence low values for the two features roundness and solidity; and (II) in a cluster (CD29^+^ cells), which was negative for CD45 and hematopoietic lineage markers and mainly composed of large cells, suggesting a mesenchymal stromal phenotype. These cells, however, displayed a low abundance of VIM. One cluster exhibited a pronounced expression of HLA-ABC, but could not be assigned to a specific cell type.

Taken together, we here provide a comprehensive representation of DTCs and the bone marrow microenvironment of neuroblastoma patients.

### Analysis of bone marrow microenvironmental changes

We then investigated the influence of DTC abundance on the bone marrow microenvironment by separately analyzing the cell composition of each individual bone marrow sample (Fig. 3a, Fig. S9).

**Figure 3:**
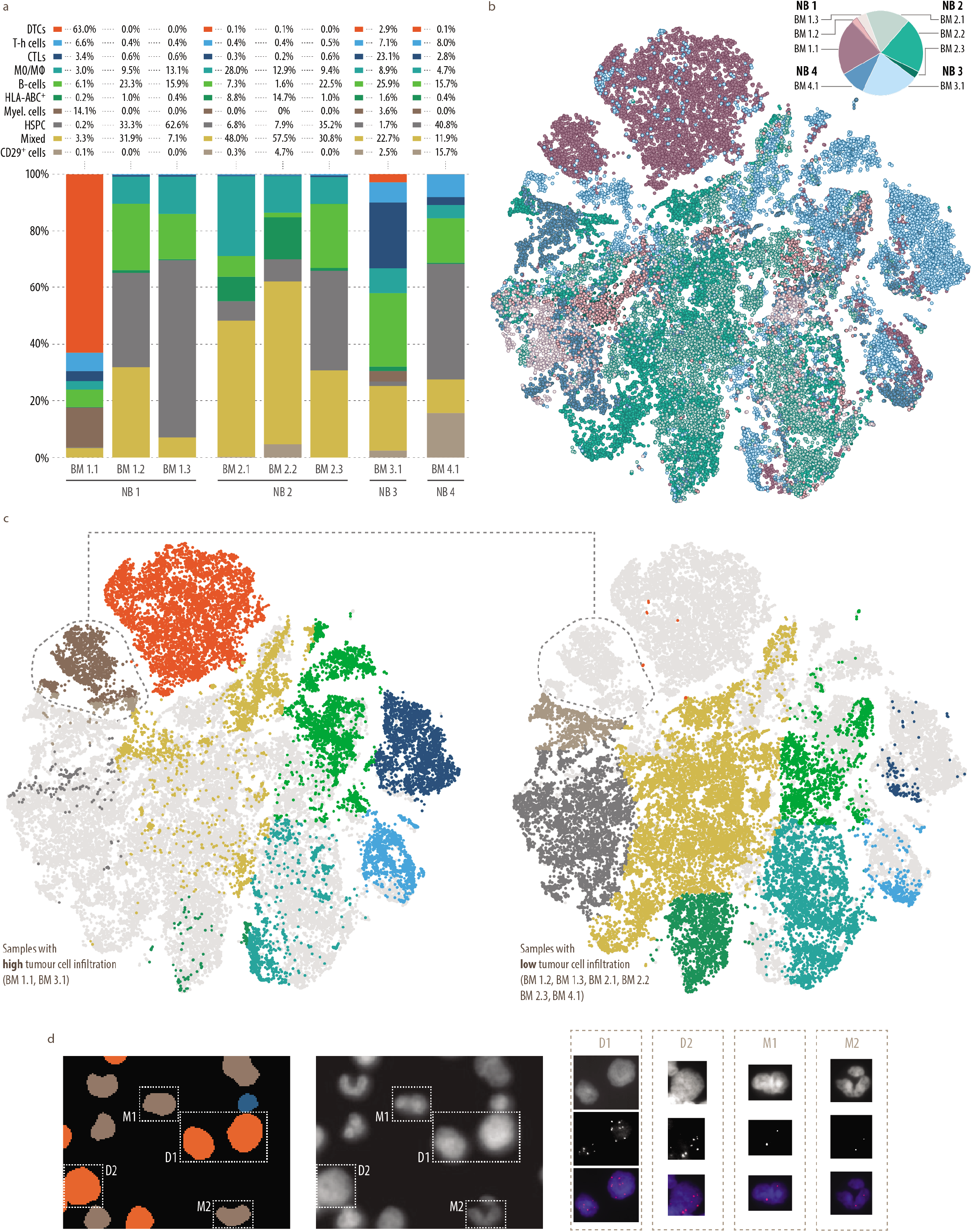
Changes in the cell composition associated with the presence of tumour cells in the bone marrow. **a,** Bar charts demonstrating the cell composition of 8 analyzed bone marrow samples. DTCs, disseminated tumor cells; Myel., myelocytes; MO/MΦ, monocytes/macrophages; Mes. cells, mesenchymal cells; HSPC, hematopoietic stem and progenitor cells; T-h cells, T-helper cells; CTLs; cytotoxic T-lymphocytes; Mixed, hematopoietic mixed cell population. **b,** A-tSNE plot of 35,700 single cells colored by sample and pie chart showing sample size. **c,** A-tSNE plot of 35,700 single cells highlighted by samples with high (top, BM 1.1, BM 3.1) and low (bottom, BM 1.2, BM 1.3, BM 2.1, BM 2.2, BM 2.3, BM 4.1) tumor cell infiltration and colored by cell type. Dimensionality reduction was performed by A-tSNE (approximated and user steerable t-distributed Stochastic Neighbor Embedding) and subsequent clustering by GMS (Gaussian Mean Shift) in Cytosplore.^50^ **d,** FISH analysis with chromosome 17q-specific probe on MELC-preprocessed sample BM 1.1, collected from a patient with 17q gain. Nucleus segmentation mask (left, large) pseudo-colored according to cell type and based on propidium iodide image (right, large) acquired during MELC. 6 copies of 17q (red) were detected on DTCs (D, orange) and 2 on myelocytes (M, brown). The 17p reference probe did not yield interpretable results due to preprocessing of the sample by MELC. Nuclei were counterstained with DAPI (blue).

Independent of the sample size (Fig. 3b), DeepFLEX detected DTCs in the same bone marrow samples as AIPF (Table 2), currently the standard method for minimal residual disease detection in routine diagnostic procedures. Although the proportion of DTCs detected by the two methods was in a similar range, absolute numbers differed, which can be explained by the higher number of markers used in DeepFLEX and the difference in sample size.

Next, we compared bone marrow samples with high and low tumor cell content (Fig. 3b and c). The proportion of hematopoietic mixed as well as stem and progenitor cells was strongly reduced in samples with a high tumor cell infiltration. Moreover, myelocytes appeared only in samples with a high DTC content.

In order to exclude that the myelocytic cluster co-expressing CD24, which is also highly expressed in DTCs, and the mesenchymal marker CD29, might contain mesenchymal-type neuroblastoma cells^19–20^, we performed interphase fluorescence in situ hybridization (iFISH) subsequent to MELC. The bone marrow sample with the highest DTC fraction and most abundant CD29^+^CD24^+^ cluster (BM 1.1) originated from a patient with a chromosome 17q gain and was therefore interrogated using a 17q-specific probe. The result (Fig. 3d) unequivocally demonstrated that cells from the myelocytic cluster do not carry supernumerary 17q signals and were therefore considered normal cells. However, these cells only appeared in the presence of DTCs in the bone marrow in our sample set. In addition, FISH analysis also confirmed the accurate classification of DTCs. We clearly detected six copies of 17q, which was in accordance with a previous FISH analysis of lymph node metastases from the same patient (NB1, Fig. S10).

Patient NB1 (Table 3) was diagnosed with a primary tumor located in the right adrenal gland, and widespread metastatic bone marrow infiltration according to abdominal magnetic resonance imaging (MRI) and the meta-iodobenzylguanidine (MIBG) scan. This was also reflected in results obtained by DeepFLEX, which detected a DTC content of 63% in the diagnostic bone marrow sample BM 1.1 (Table 2, Fig. 3a). Upon induction chemotherapy, the patient showed a good local response and no evidence of tumor cells in the bone marrow, which was in accordance with our results (BM1.2), (Table 2, Fig. 3a). Therapy response also coincided with the expansion of hematopoietic mixed as well as stem and progenitor cells indicating hematopoietic restoration.

In summary, we observed that dissemination of neuroblastoma tumor cells into the bone marrow as well as response to therapy was associated with changes in the bone marrow microenvironment, specifically alterations of the myelocyte cell and hematopoietic mixed and stem and progenitor cell compartments.

### Heterogeneity of disseminated tumor cells and FAIM2 as a novel complementary marker

Though neuroblastoma tumor heterogeneity has been investigated at the primary site on a single-cell level^19,20^, a respective characterization in the metastatic bone marrow was still missing. To assess tumor heterogeneity of bone marrow metastatic cells, we therefore performed hierarchical clustering (see Methods) on the single-cell data of our DTC cluster using only the DTC markers from our 20-plex MELC panel.

We obtained a clustermap with 30 DTC sub-clusters showing the heterogeneous expression of markers expressed by DTCs (Fig. 4a), which was also reflected by representative gallery images of individual DTC phenotypes (Fig. 4b). Notably, half of all DTCs belonged to sub-cluster 19, which represented a dominant phenotype in the two highly tumor-infiltrated bone marrow samples BM 1.1 and BM 3.1 (Fig. 4c). While DTCs showed a predominantly round nuclear shape, cellular and nuclear size contributed to the fractionation of DTCs into distinct sub-clusters and varied between different phenotypes, e.g. 17 and 30, which were composed of mainly large and small cells, respectively. Sub-cluster 18 and 20 displayed a high expression of FAIM2, an inhibitory protein in the Fas-apoptotic pathway of tumor cells^53,54^, which was proposed as a tumor marker in small cell lung^55^ and breast cancer^56^. FAIM2 is known to be primarily expressed in neurons^54,57^, which was in accordance with our bulk proteomics data (Fig. S3a). In our RNA-seq datasets, FAIM2 transcription was significantly higher in tumor cells than in bone marrow-derived MNCs (Fig. S3b). Moreover, FAIM2 transcription was enriched in primary tumor cells without *MYCN* amplification as compared to those with *MYCN* amplification (Fig. S3B), thus supporting previous findings^58^. However, we did not observe a differential expression between these two classes in DTCs. Interestingly, in our single-cell analysis of eight neuroblastoma bone marrow samples, FAIM2 was expressed only in a subset of DTCs (Fig. 4a and b, Fig. S11). As other markers, that were found to be expressed by DTCs, FAIM2 was NOT exclusive to neuroblastoma cells, but was also found on other cell types in the bone marrow (Fig. S8 and S11). To assess the correlation of FAIM2 and other DTC markers, we plotted the DTC marker abundances for all cells of the DTC clusters (Fig. 4d, Fig. S12). This corroborated the observation, that only a subset of DTCs exhibit a high expression of FAIM2 along with low or intermediate abundances of the other DTC markers, while the latter were mostly co-expressed by DTCs.

**Figure 4:**
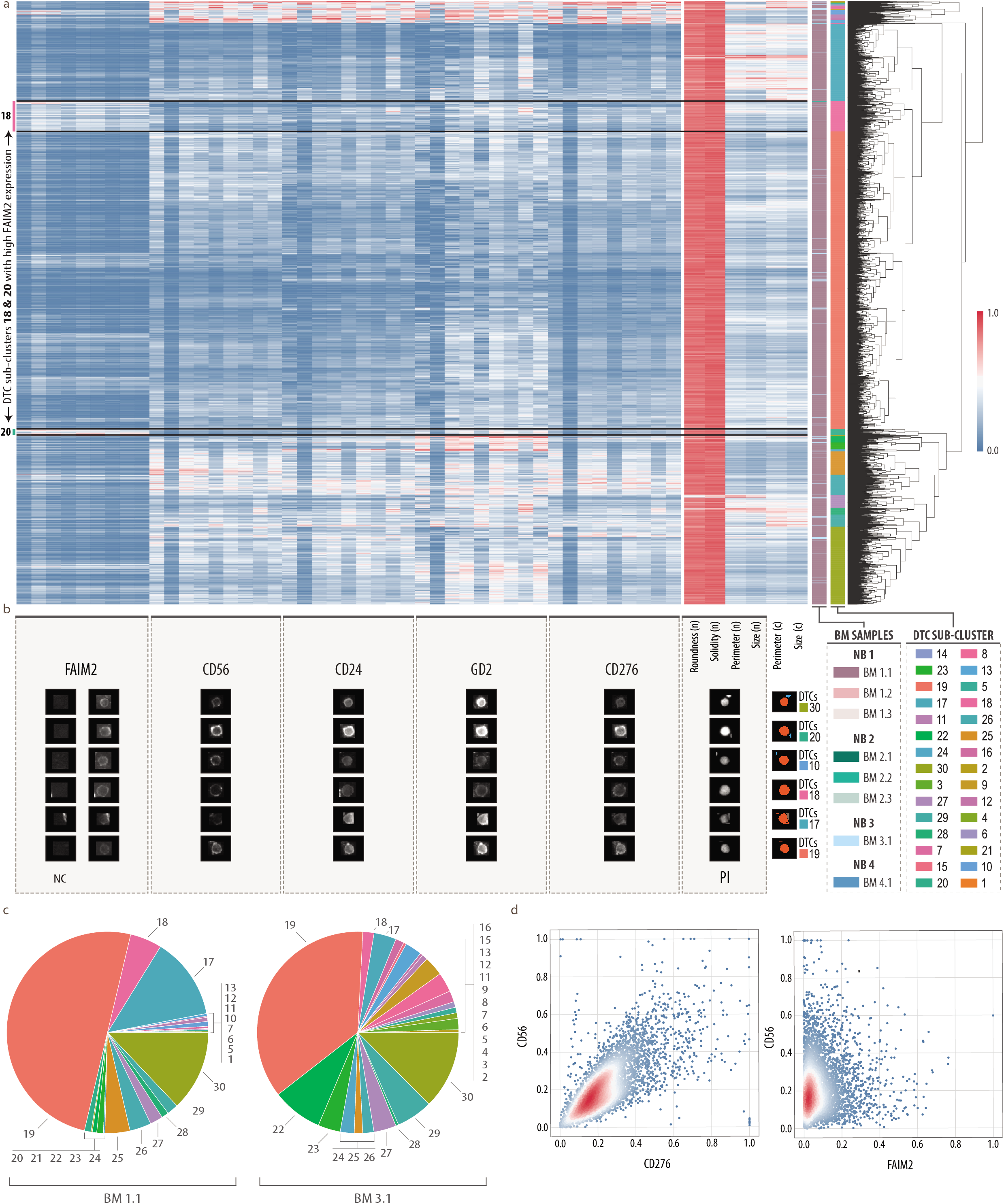
Characterization of DTC heterogeneity and qualification of FAIM2 as a novel complementary biomarker. **a,** Clustermap (hierarchical clustering by Voorhees^80^) showing normalized single-cell feature values of DTCs. n, nucleus; c, cell. 9 columns per marker represent, from right to left, mean intensity, total intensity and mean of the highest 20% of pixel values in the (I) nucleus, (II) cell and (III) cytoplasm/membrane. Color bar on the right shows 30 sub-clusters. Color bar on the left shows corresponding bone marrow sample. **b,** Representative gallery images of 6 selected cells from different DTC sub-clusters reflecting DTC heterogeneity. For FAIM2 we introduced negative controls (NC) to be used as background threshold levels during data processing. Hence, for this biomarker, the ratio between right column and left column (NC) represents the true signal. **c,** Proportion of 30 DTC sub-clusters in highly tumor-infiltrated bone marrow samples (BM 1.1, BM 3.1). **d,** Scatter plots showing correlations of DTC marker CD276 versus CD56 (left), and FAIM2 versus CD56 (right) for all cells of the DTC cluster. Mean of the highest 20% of pixel values in the cytoplasm/membrane was used as measure for marker abundance.

Here we present a first exploratory survey of DTCs in bone marrow metastases on a single-cell level highlighting a hitherto unappreciated diversity pointing toward multiple distinct subclasses of DTCs. We show that FAIM2 marks a subset of DTCs and can serve as a complementary biomarker for capturing DTC heterogeneity in neuroblastoma.

## DISCUSSION

While the bone marrow attracts tumor cells in numerous solid cancer entities leading to poor outcome in affected patients, comprehensive analyses of bone marrow metastases have not been performed on a single cell level. We here set out to capture tumor heterogeneity and unravel microenvironmental changes in a solid cancer with bone marrow involvement.

To this end, we constructed an atlas of DTCs and their microenvironment in the metastatic bone marrow niche by multiplex imaging of eight human neuroblastoma bone marrow samples and subsequent image analysis by our newly-developed pipeline DeepFLEX. Our results revealed vast diversity among DTCs and suggest that FAIM2 can act as a complementary marker to capture DTC heterogeneity. The presented findings indicate that malignant bone marrow infiltration and response to cancer therapy might be associated with changes in the bone marrow microenvironment, warranting deeper investigations of spatio-temporal dynamics at the single-cell level and of their clinical relevance.

The bone marrow, as part of the immune system, constitutes a niche comprised of multiple immune cell subpopulations^59^, shown to be involved in cancer progression.^60^ As a key regulator of hematopoietic and mesenchymal stem cell function, the niche may facilitate quiescence and drug-resistance^61^, impairing current therapeutic approaches. Single-cell multi-modal analysis of healthy human bone marrow recently identified the major bone marrow mononuclear populations.^51^ However, the single-cell atlas of malignant human bone marrow has so far only been described in leukemia^62,63^, where the bone marrow is not considered a metastatic, but rather an originating site. Herein, we provide first insights into the single-cell landscape of human bone marrow metastases including variations among DTCs as well as cells of the mesenchymal and hematopoietic compartment.

Among DTCs, we showed a high level of diversity reflected by heterogeneous cell morphologies as well as protein expression profiles and fractionation into phenotypically diverse DTC sub-clusters. Notably, half of the cells belonged to one major DTC sub-cluster, which represented a dominant phenotype in both of the two highly tumor-infiltrated bone marrow samples. This phenotype dominance was also observed in a previous study^64^ on breast cancer, where in almost half of the analyzed cohort, 50% of all tumor cells belonged to a single tumor cluster, and might reflect clonal expansion, intrinsic plasticity or result from tumor-microenvironment interaction.

A subset of DTCs exhibited a high expression of FAIM2, an inhibitory protein in the Fas-apoptotic pathway, which we included into the 20-plex panel upon data mining of a previously generated neuroblastoma RNA-seq and proteomics dataset. FAIM2 was described as a therapeutic target in small cell lung cancer^55^ and as a predictive marker of poor outcome in breast cancer patients.^56^ Herein, we propose FAIM2 as a complementary marker to depict a broader spectrum of DTC heterogeneity, as it marked a subpopulation of DTCs and showed a lower correlation with the other analyzed DTC markers than the latter with each other. DCLK1, a cancer stem cell marker^65^, was another candidate of high interest, whose distribution is yet to be assessed in bone marrow metastases. A deeper investigation of DTC sub-classes in larger patient cohorts may yield targets for neuroblastoma therapy.

Neuroblastoma heterogeneity has been investigated before by scRNA-seq of primary tumor samples, most of which were shown to be composed of adrenergic and mesenchymal type neuroblastoma cells, the latter suggested to be more resistant to chemotherapy.^20^ Also, we have previously shown that mesenchymal characteristics can be adopted upon therapy-induced tumor cell senescence.^66,67^ Yet, in the investigated bone marrow samples, collected at different time points in the disease course, we did not detect neuroblastoma cells of a mesenchymal phenotype based on the expression of the mesenchymal markers, CD29 and Vimentin. This might be explained by the limited sample as well as panel size and the fact that mesenchymal type neuroblastoma cell identity has previously been defined by master transcription factors active in gene regulatory networks. Thus, future research will require the identification of robust imaging-based markers to reliably assign neuroblastoma cells to these two classes.

Within the bone marrow microenvironment, we observed alterations in the hematopoietic and mesenchymal cell compartment with respect to the level of tumor cell infiltration indicating that DTCs shape the metastatic niche, albeit based on a very limited cohort size. In support of this notion, leukemia cells are likewise known to reprogram the bone marrow niche in order to instigate changes that promote their progression.^68^ Interestingly, we identified considerably fewer progenitor and other immature hematopoietic cells (distributed among 2 clusters, i.e. hematopoietic mixed and stem and progenitor cells), in highly tumor infiltrated samples, which solidifies previous findings, suggesting that tumor invasion reduces the support for primitive hematopoietic stem and progenitor cells in the metastatic niche.^69^ In addition, it is widely accepted that cytotoxic therapy leads to bone marrow perturbation resulting in myelosupression^70^, which might also be responsible for the observed depletion of immature, lineage negative hematopoietic cells. Furthermore, we only found myelocytes in samples with a high DTC infiltration, which may be attributable to the immunosuppressive and tumor-promoting functions of cells from the myeloid origin as well as their role in inflammation.^71^ Our findings were based on single-cell analyses of MELC multiplex imaging data, enabled by the pipeline DeepFLEX, which we developed based on the integration of methods for image processing^45,46^, segmentation^47,48^, normalization^49^ and single-cell analysis.^50^ DeepFLEX tackles confounding factors of targeted imaging technologies such as unspecific binding and autofluorescence and combines deep-learning based cell and nucleus segmentation, which allow accurate single-cell assessment. In addition to the code, we provide the complete multiplex image dataset of all samples used in this study. The demonstrated application of DeepFLEX on MELC imaging data serves as a blueprint for further single-cell analyses by multiplex imaging methods beyond MELC, facing similar challenges.

In conclusion, this study offers a first view into the single-cell landscape of human bone marrow metastases and might motivate further investigations in other solid cancers with bone marrow involvement. Moreover, our findings represent a valuable source of information for the design of therapeutic approaches depending on the distribution of target molecules on cancer cells such as immunotherapies^72^, and can hence contribute towards better patient stratification in neuroblastoma.

## DATA AND CODE AVAILABILITY

The RNA-seq dataset used for datamining is available for download on the GEO data repository under accession number GSE94035.

Supplementary data (.csv and .doc files) holding manually curated information retrieved from protein databases and literature search can be found on https://cloud.stanna.at/sharing/iyorsYWzp.

The mass spectrometry proteomics data has been deposited to the ProteomeXchange Consortium (proteomecentral.proteomexchange.org) via the PRIDE partner repository (ebi.ac.uk/pride/, PMID: 24727771) with the dataset identifier PXD018267.

Python code for the DeepFLEX pipeline is available on github.com/perlfloccri/DeepFLEX. A compiled release with all necessary dependencies pre-installed is available from dockerhub URL https://hub.docker.com/repository/docker/imageprocessing29092020/deepflex.

The MELC multiplex imaging data of our neuroblastoma cohort is available at https://cloud.stanna.at/sharing/qiN0u9QPO.

## ACKNOWLEDGEMENTS

This work was facilitated by the Austrian Research Promotion Agency FFG (Project Visiomics, grant no. 10959423 to S.T.M.), Austrian Science fund FWF (Project Liquidhope within the ERA-Net/Transcan-2 program, grant no. I 4162 to S.T.M.) and by private donations to St. Anna Children’s Cancer Research Institute (Vienna, Austria). We thank all patients and parents, who participated in the study. Prof Handgretinger, University of Tübingen kindly provided the anti-GD2 antibody. For excellent technical support, we thank Andrea Ziegler, Bettina Brunner-Herglotz, Anja Zimmel and Maria Berneder.

## AUTHOR CONTRIBUTIONS

D.L., F. K., I.M.A., P.F.A., C.O. and S.T.M conceived the study. M.U., M.B. and R.L. provided clinical samples. F.R., A.B. and C.G. generated the transcriptomic and proteomic dataset, respectively. D.L. developed the antibody panel and prepared samples for MELC. F.R. assisted in the interpretation of transcriptomic data and in data visualization as well as illustration. C.O. and M.K. carried out MELC. D.L., F.K. and F.M. designed and developed DeepFLEX. F.H. contributed data processing methods. D.L. and S.T.M. analyzed and interpreted the single cell data. D.L., F.K. and S.T.M. wrote the manuscript with input from all authors. D.L., F.K., M.K., F.M., F.R., F.H., M.B., A.B., M.U., I.M.A., P.F.A., C.G., R.L., C.O. and S.T.M. revised the manuscript.

## CONFLICTS OF INTEREST

The authors declare no conflicts of interest.

## METHODS

### Patients and cell lines

The collection and research use of human tumor specimen was conducted according to the guidelines of the Council for International Organizations of Medical Sciences (CIOMS) and World Health Organization (WHO) and has been approved by the local ethics committees of the Medical University of Vienna (EK1216/2018, EK2220/2016).

#### Neuroblastoma cell lines

Five patient-derived neuroblastoma cell lines were used for validation of biomarkers and identification of the best sample preparation conditions. STA-NB-2, −4 and −10 have been established from primary tumors, and STA-NB-8 and STA-NB-12 from DTCs of bone marrow aspirates. INSS (International Neuroblastoma Staging System) and *MYCN* amplification status for all neuroblastoma cell lines were described previously^73^ and are listed in Table S2. Cells were maintained in RPMI1640-Glutamax-I (GIBCO) supplemented with 1% Pen/Strep (GIBCO), 10% FCS (PAA Laboratories), 1 mM sodium pyruvate (PAN Biotech) and 25 mM HEPES (PAN Biotech). All neuroblastoma cell lines were cultivated at 37°C and 5% CO2.

#### Bone marrow aspirates

Bilateral bone marrow aspirates were collected according to the SIOPEN/HR-NBL-1 study protocol or standard of care during routine diagnostics at initial diagnosis and at clinical response evaluation time points. Samples were shipped at room temperature within 4 hours or at 4°C within 24 hours. Bone marrow-derived MNCs were isolated by density gradient centrifugation (LymphoprepTM, AXIS-SHIELD PoC AS).

For the validation of antibodies, neuroblastoma cell lines were mixed with tumor-free bone marrow-derived MNCs to obtain a tumor cell suspension of 5% neuroblastoma cell line in bone marrow-derived MNCs.

For single-cell analysis, eight bone marrow aspirates (Table 2) were collected at different time points along the therapy protocol from four neuroblastoma patients with metastatic (INRG stage M or Ms) disease (Table 3).

#### Peripheral blood-derived MNCs

Left over samples of peripheral blood from routine diagnostics was collected and peripheral blood-derived MNCs were isolated, washed and counted as described for bone marrow-derived MNCs. Neuroblastoma cell lines were spiked into peripheral blood-derived MNCs to obtain a tumor cell content of 5%. The cell mixture was cultivated in the presence 0.125% (v/v) anti-CD3/CD28 beads (Thermo Fisher Scientific) and 1% (v/v) IFNγ (Peprotech) for 5 days.

### Biomarker identification by data mining

DTC-associated biomarkers were identified based on data mining of previously generated RNA-seq datasets and proteomics data, and guided by public databases. The following prioritization scheme was employed:

#### Differential gene expression analysis

RNA-Seq data (GEO repository available under accession number GSE94035) of primary tumors (n=16), enriched bone marrow-derived diagnostic (n=22) and relapse DTCs (n=20), and the corresponding bone marrow-derived MNCs (n=28) of in total 53 stage M neuroblastoma patients was processed as previously described^18^ and used for the identification of potential DTC biomarkers. Genes with significantly higher (DEseq2^74^, FDR-adjusted p ≤ 0.001, log2FC ≥ 4) transcript levels (FC, fold change) in DTCs as compared to bone marrow-derived MNCs were selected (n=1,594) and further filtered for those with an equal or higher transcription (DEseq2, FDR-adjusted p : 0.01 ÷ 0.7, log2FC ≥ 0) at the time point of relapse as compared to diagnosis (n=921).

#### Protein databases and literature search

The remaining genes were manually annotated with the cellular location of the encoded protein according to protein databases UniProt^75^ and The Human Protein Atlas^76^, and only proteins localized on the cell membrane by at least one database were further considered (n=134, see Data and code availability).

Detailed literature search using the search terms [neuroblastoma], [tumor] and [metastasis] was carried in the PubMed database (pubmed.ncbi.nlm.nih.gov) resulting in 99 candidates (see Data and code availability), from which five (TAG1, DCLK1, FAIM2, PRAME and TACC2) were selected based on detailed examination of available literature and commercial availability of respective antibodies.

#### Proteomics data

Proteomics data of eight peripheral-nerve-associated fibroblasts, 3 in-house established patient-derived neuroblastoma cell lines (STA-NB-10, STA-NB-2, STA-NB-7) and 6 corresponding neuroblastoma primary tumors was previously generated^77^ and is available on the ProteomeXchange Consortium (proteomecentral.proteomexchange.org) with the dataset identifier PXD018267. The proteomics dataset was used to confirm the expression of the five candidates selected above as well as seven other biomarkers (CD56, NCAM-L1, PD-L1, VIM, PROM1, B7-H3 and PD1), which were added based on their relevance in neuroblastoma as previously reported. ^20,32,33,34,35^

### Biomarker validation

#### Cytospin slide preparation

50,000 cells (cell lines) or 250,000 cells (spike-in and patients samples) were applied onto poly-L-lysine hydrobromide (PLL) (Sigma Aldrich) -coated microscope cover glasses (24×60 mm, Assistent) using filter paper (4 ml, CytoSepTM) and funnel chamber (4 ml, CytoSepTM) of a Hettich cyto-centrifuge (Hettich). Three different centrifugation and fixation methods were tested in the present study (Table S1, Fig. S1b, Fig. S2a). The optimized protocol for processing patient samples is detailed in Table S1 and involves PFA (paraformaldehyde) followed by acetone (AC) fixation (PFA-AC). Chemicals used for fixation, acetone and 4% PFA, were ordered from Carl Roth GmbH. Slides were dried for 2 min after fixation and stored at −80°C until further analyses.

#### IF Staining

Antibodies (Table S4) were diluted in 2% BSA/PBS. Slides were incubated with primary antibody solutions for 1 h at room temperature, washed in PBS twice followed by secondary antibodies for 1 h at room temperature. After washing, slides were incubated with the nuclear stain DAPI (2 μg/ml) for 2 min and covered with antifade medium Vectashield (Vector Laboratories).

#### Validation procedure

DTC-related biomarkers were validated based on an intuitive validation procedure (Fig. S1a). First, IF-staining of individual biomarkers was performed on neuroblastoma cell lines prepared with the AC and PFA based protocol (Table S1, Fig. S1b). Thereafter, slides were assessed visually using a Zeiss Axioplan two microscope in five criteria (nuclear morphology, background noise, cell debris, staining intensity, staining quality) to evaluate the impact of the respective sample preparation protocol (AC or PFA) on cell morphology and antigenicity. Scores from one to five were assigned to each criterion with five corresponding to the best result. Accordingly, the maximum score for one slide was 25. Overall scores for all five neuroblastoma cell lines, incubated with the corresponding antibody, were summed up for each fixation method separately, and a mean score was calculated as a qualitative metric (Table S3). Antibodies with a mean score below 13 for both AC and PFA based fixation were considered invalid and not further validated. For all other antibodies, images of the slides prepared with the better sample preparation protocol (higher mean score) were acquired using the automated scanning system, Metafer 4 (software version V3.11.8 WK, Metasystems) and 63x magnification (Fig. S1c).

Antibodies that were successfully validated on neuroblastoma cell lines, were additionally tested on two cytospin samples of neuroblastoma cell lines and bone marrow-derived MNCs or peripheral blood-derived MNCs (for validation of PD-L1, PD-1) prepared with the PFA-AC protocol (Table S1, Fig. S2a). Slides were then visually inspected and imaged automatically as above (Fig. S2b).

For sequential IF-staining by MELC, antibodies, which passed the validation procedure, were combined with already validated antibodies specific to bone marrow hematopoietic and mesenchymal cells (Table S4). Staining sequence and panel were refined in several pilot MELC rounds and finally resulted in a 20-plex biomarker panel (Table 1, Fig. 1e and f).

### Multi Epitope Ligand Cartography

MELC was employed for multiplex IF-staining of the herein established 20-plex antibody panel, as described.^30^

Briefly, MELC is based on repetitive cycles of antibody staining and photobleaching. After system start, four field of views are selected and calibration (brightfield and darkframe) images are acquired. Prior to every staining and photobleaching cycle with the acquisition of the corresponding fluorescence tag and post-bleaching image, the slide is washed with PBS and a phase contrast image is taken.

Camera (ApogeeKX4,Apogee Instruments) and light source maintain the same position; the motor-controlled xy stage of the inverted fluorescence microscope (Leica DMIRE2, Leica Microsystems; x20 air lens; numerical aperture, 0.7) moves in between field of views. Images with a resolution of 2018 x 2018 pixels are acquired, with one pixel corresponding to 0.45 μm at a 20x magnification. Thus the whole image covers a field of view covering 908.1 x 908.1 μm.

Additionally, negative control secondary antibodies were implemented, which were applied to the sample prior to indirect staining of the respective primary antibody.

### Interphase fluorescence in situ hybridization (iFISH)

The MELC pre-processed sample BM 1.1 was fixed in 4% paraformaldehyde at 4°C overnight for subsequent analysis by iFISH. iFISH was performed as previously described.^78^ Predigestion of cells was carried out in 0.005% pepsin in 0.01 NHCL for 25 min. Since the sample originated from a patient with a chromosome 17q gain, a labeled 17q-specific probe (XL Iso (17q), Metasystems probes) was used. Denaturation was performed at 80°C. Nuclei were counterstained with nuclear stain DAPI (2 μg/ml) for 2 min and covered with antifade medium Vectashield (Vector Laboratories). Slides were imaged with the Zeiss Axioplan 2 microscope and the ISIS software (version 5.7.4, Metasystems).

### DeepFLEX

Main parameters used by methods integrated in the pipeline are listed below. Further parameters are detailed in Table S8.

#### Image processing

Images were registered, as previously described^45^ (Fig. S4a). Then, flat field correction using brightfield and darkframe calibration images was performed to eradicate gross variations in illumination (Fig. S4b). Accumulative background noise caused by residual post-bleaching signals was eliminated by subtracting post-bleaching images from successive fluorescence tag images (Fig. S4c). To reduce vignetting (reduction of image brightness toward periphery compared to image center), intensity distributions were corrected using regularized energy minimization on the set of all fluorescence tag images from our eight samples via CIDRE^46^ (github.com/smithk/cidre); (Fig. S4d).

#### Segmentation

For accurate nuclei and cell segmentation, annotated datasets^48^ of propidium iodide or phase contrast images were created, respectively. We trained the deep learning architecture Mask R-CNN for instance-aware segmentation, as previously described.^47^ Briefly, after augmenting the training dataset with automatically generated artificial images, we used image tiling and rescaling to segment MELC images in order to make them compatible with the input size (256×256 pixels) of the trained Mask R-CNN. The fluorescence tag image (nuclear stain propidium iodide) was segmented into a labeled nucleus mask (Fig. S4f), while phase contrast images (which are acquired prior to each IF staining) were segmented into labeled cell masks (Fig. S4e) for each of the 20 markers. Inferred objects were only counted as cells if they were reproduced in all of the 20 cell and the nucleus mask (Fig. S4g). We furthermore removed cells affected by image artifacts or located in poorly illuminated image corners by user-guided region selection (Fig. S4h).

#### Feature extraction

The segmentation masks were used as a reference to generate multi-channel single-cell images (Fig. S5a), based on which intensity and morphological features were extracted (Fig. S5b, Table S5). The morphology of the cell nucleus was described by the features size, perimeter, roundness and solidity. To describe the morphology of the cell, the features size and perimeter were extracted. We used three intensity features to quantitate marker abundance: mean intensity, total intensity and mean of the top 20% intensities (less dependent on cell size). Intensity was measured per cell, per nucleus, and per cell cytoplasm and membrane (= cell - nucleus). After feature extraction, cells with a larger nucleus than cell segmentation mask were excluded.

#### Normalization

To eliminate unspecific staining caused by secondary antibodies and to increase the signal-to-noise ratio, features extracted from the second secondary antibody were divided by features extracted from the negative control secondary antibody (Fig. S5c).

To further remove unspecific binding of primary antibodies and batch variation in staining intensity, autofluorescence, and illumination, we applied RESTORE^49^ (gitlab.com/Chang_Lab/cycif_int_norm) to predict a background level (threshold separating signal and noise/background) for each marker in each image based on a mutually exclusive counterpart. Mutually exclusive marker pairs (Table S6) were selected based on biological knowledge and a data-driven approach using singular value decomposition, as described.^49^ Background levels were then inferred for each field of view and each intensity feature, separately. Guided by generated scatter plots (Fig. S5d, top), we selected the background level predicted by sparse subspace clustering (*σ* = 0). If no positive signals were present in the analyzed field of view for a certain marker by visual inspection, the respective background level was set to the maximum intensity value.

Subsequently, all values below the background level were randomly set within a range between 0 and 0.02, while all values exceeding the background level (corresponding to signals) were linearly scaled to a range between 0.02 and 1. Thereby, influence of background variation on the subsequently applied single-cell analysis was eliminated, while foreground signals were stretched to a larger dynamic range.

Morphological features were linearly scaled between 0 and 1. Upon RESTORE normalization and scaling, batch effects were successfully removed (Fig. S5d, bottom).

#### Single-cell analysis

Normalized features were converted into an FCS file format and loaded into Cytosplore^50^ (cytosplore.org, version 2.3.1), an interactive tool providing methods for single-cell analysis. A-tSNE^79^ (approximated and user steerable t-distributed Stochastic Neighbor Embedding, *perplexity* = 30) and subsequent clustering by GMS^80^ (Gaussian Mean Shift, *σ* = 45) clustering was computed on the complete single-cell dataset of eight bone marrow samples and resulted in 10 clusters, which were exported as FCS files together with the CSV file of the corresponding heatmap (Fig. 2b and c). The latter were imported into python to allow further quantitative and explorative analysis (Fig. 2d, Fig. 3a-d, Fig. 4a-d, Fig S7-S9) with the python (version 3.7) data visualization library seaborn (seaborn.pydata.org, version 0.10.rc0), (Fig. S5e). Hierarchical clustering was performed using the complete-link / Voorhees algorithm^80^ provided by seaborn.

## Notes

### Competing Interest Statement

The authors have declared no competing interest.

